# The role of startle fluctuation and non-response startle reflex in tracking amygdala dynamics

**DOI:** 10.1101/2020.01.12.903526

**Authors:** Mengting Liu, Rachel C. Amey, Adam Magerman, Matthew Scott, Chad E. Forbes

## Abstract

The startle reflex is considered a sensitive defensive reaction to potential threats that manifests as a unique eye blink-like pattern in the EMG. Eye blink EMG has a bell-shaped potential when startle probes are elicited, that strongly corresponds to amygdala activity. Considering how amygdala activity fluctuates over time in response to emotional and self-threatening stimuli, observing startle eyeblink size fluctuation over time could provide a cost-effective, convenient, and less resource intensive means for examining amygdala activity over time. Yet based on current standards in the literature, often startle evoked eye blink EMG values do not exhibit activity 3SDs from the mean eyeblink response, thus these trials are typically excluded from startle analyses. It stands to reason, however, that these trials may still index amygdala activity in a meaningful way. Through investigating the association between startle eyeblink amplitude, corresponding ERP amplitude, and underlying neural activity, the current study provides evidence that startle amplitudes exhibit a linear relationship with proxies for amygdala activity, e.g., N100 amplitudes and regions heavily interconnected with the amygdala. Specifically, the startle reflex correlates to large amount of brain regions in N100 time window in addition to the N100 amplitude. Thus, both valid and otherwise traditionally non-valid startle reflex responses appear to index amygdala activity and should be included accordingly. This approach could help salvage large amounts of meaningful data traditionally excluded from studies interested in amygdala responses to various stimuli over time.

---

The startle reflex is considered a sensitive defensive reaction to potential threats that manifests as a unique eye blink-like pattern in the EMG. Typically, these blink responses have a normal bell-shaped curve above baseline activity, and are often elicited by presenting loud acoustic noises via headphones or puffs of air to the eyeball (Lissek, et al, 2005; Yeomans & Frankland 1995). Research has demonstrated that the magnitude of the startle amplitude is directly related to amygdala activity (LeDoux, 1996; Alessandro, et al. 1996). Conventionally, startle eyeblink sizes are measured by averaging startle eyeblink amplitude across all trials in a single condition. However, as past amygdala research has revealed a more dynamic, non-linear pattern of activity towards emotional stimuli (Phillips et al., 2001), tracking single trial startle eyeblink amplitudes over time seems particularly relevant and worthwhile as they may ultimately reflect more nuanced amygdala patterns.

Yet, often individuals in startle experiments do not always exhibit a startle response that reaches criteria, especially after habituation, despite reporting increased arousal (Bradley, Lang, Cuthbert, 1993). Past startle based research has dealt with these trials in two ways specifically, either by excluding these eye blink responses (Blumenthol, 2005), or replacing these trials with an average startle amplitude (e.g., Bradley, Codispoti & Lang, 2006; Nelson, Hajcak & Shankman, 2015; Jiang et al., 2016). Are these excluded trials devoid of amygdala activity? In relation to current conceptualizations of the amygdala they are likely not. To date, little is known about what exact neural processes are reflected in both above and below threshold startle single trials. Understanding whether below threshold startle trials are meaningful could provide a means to observe amygdala fluctuations over time in a more cost-effective, convenient and less resource intensive way than the only other option currently available-functional magnetic resonance imaging (fMRI).

In this study, the properties of both above and below threshold startle responses were examined in conjunction with their underlying neural activity. Emphasis was placed on understanding what startle amplitudes fluctuation reflect with respect to specific brain regions and event related potentials (ERPs), especially for non-response trials, to determine if these trials provided meaningful data with respect to amygdala activity. Results suggest that non-response startle trials exhibit identical underlying neural activity as valid startle trials, suggesting meaningful data specific to the amygdala is evident in all startle trials, regardless of whether they are traditionally deemed valid. Moreover, including all trials replicated past research that has demonstrated a quadratic relationship between amygdala fluctuation and responses to threatening compared to neutral stimuli over time.

## Startle Probe Methodology to Date

Startle responses are captured and measured by eyeblink patterns in EMG recordings. Usually trials with a peak value larger than a predefined threshold (three standard deviations higher than mean eyeblink responses) are considered valid response trials (Brinkworth & Tucker, 2003). Trials that do not reach criteria are considered invalid, or non-response trials (Bluementrol, 2005). The size of a valid startle eyeblink has not been officially defined regarding magnitude and amplitude. When discussing a single trial, magnitude and amplitude are analogous. However, when discussing the average of multiple trials, magnitude and amplitude take on different meanings (Blumenthal, 2005). Magnitude has been utilized when the average of multiple trials includes trials with zero-values (Quevedo, et al., 2010; Hillman, Hsiao-Wecksler & Rosengren, 2005; Rhudy, et.cl., 2008), while amplitude has been used when the average is computed with invalid trials excluded (Klorman, et.cl., 2002; Engelmann, Gewirtz & Cuthbert, 2011; Herbert, et.cl., 2006). Other subjective approaches have been used to determine valid startle responses, for example, deflection and return to baseline in certain time intervals after stimuli (Klorman, et al., 2003), minimum duration, (e.g. Quevedo, et al., 2010), and log transforming the data (Forbes et al., 2005; Grad & Kring, 2007). To date alternative methodology for determining starting criteria has been proposed (Bluementrol, 2005), thus variations within startle results are inevitable from study to study.

Reporting non-response trials varies greatly across studies in terms of how many trials among total trials are labeled as non-response trials, from fewer than 5% (e.g. Meyer et al., 2014) to more than 90% (e.g. Klorman, et al., 2003; Quevedo, et al., 2010). Given factors like habituation to startle probes, or subsections of the population that fail to elicit startle responses at all, many if not most of startle trials are often excluded from analyses. However, given past work on the amygdala, these varying amplitudes may still provide meaningful information that arguably should not be discarded. Thus, understanding the meaning of non-response trials and providing a method to include them would be a critical advance for studies employing this approach.

## Neural ERP and Region markers can provide insight into the startle response

ERPs are often utilized to examine neural activity elicited in response to startle probes. Thus indexing ERP magnitudes is one way to examine whether there are differences between above and below threshold startle responses and whether below threshold responses are meaningful with respect to amygdala activity. Past research clearly indicates that the amygdala plays an integral role in emotional and attentional allocation processes. Thus two ERPs most often utilized to index the startle response are the N100, or the negative deflection that occurs around 100ms after the onset of startle probes, and the P300; the positive deflection that occurs around 300 ms after the onset of startle probes (Putnam & Roth, 1990; Roth, Dorato, & Kopell, 1984; Sugawara, Sadeghpour, Traversay, & Ornitz, 1994; Jiang et al., 2015).

The N100 is utilized as an index of startle responses given its clear role in early sensory processing (Olofsson, et.cl. 2009); it has also been shown to be sensitive to arousal and the physical intensity of stimuli (Naatanen & Picton, 1987). Similarly, the P300 appears to be a clear reflection of somewhat later stage attentional reorientation and encoding processes (Polich, 2007); given this, not surprisingly it too has been shown to fluctuate in relation to emotionally arousing stimuli (Cuthbert et al, 1998). The P200 may also provide insight in to discrepancies between above and below threshold startle responses given its role in downstream emotional and cognitive processes (Somme, Molen & Pascalis, 2016). Moreover, it has been implicated as a reliable measure of sensory motor gating, making it an additional measure of attentional allocation (Boutros, et.cl., 1999; Pascalis, Cozzuto & Russo, 2013).

Understanding how these components relate to startle responses could provide insight into what exactly non-response trials mean. For example, if startle trials that are above or below threshold have similar N100 waveforms then it can be suggested that amygdala responses to arousing stimuli are similar between the two trial types, at least with respect to early sensory and emotional processes. Similarly, if below and above threshold trials have similar P200 and P300 waveforms, it’s possible that these two trial types contain similar latter stage attentional and encoding processes with respect to the amygdala. Ultimately if ERP responses do not differentiate between these two types of startle trials, it could be posited that below threshold startle trials are just as meaningful as above threshold trials despite blink responses for these trials having smaller amplitudes and magnitudes.

In addition, given significant variability often found among ERPs, examining the shape of the N100, P200, and P300 curves, and isolating the neural generators of these waveforms could provide meaningful insight into whether above and below threshold trials are related. For example, although the ERP waveforms may differentiate between above and below threshold trials, if activation within the neural generator of the N100 time frame, e.g., within auditory cortex, is similar (Swick et al., 1994), it’s possible that above and below threshold responses may be analogous during this time period and similarly reflect amygdala activity accordingly. Likewise, if activation within the neural generators of the P200 or P300, e.g., within left superior temporal gyrus and parietotemporo-occipital cortex areas for P200 (Qiu et al., 2008) and parietal, inferior temporal, insula and frontal areas for P300 (Bledowski et al., 2004), were different among above and below threshold trials it stands to reason that below threshold responses do not reflect amygdala activity compared to above threshold responses within this time period.

## Study Overview

In this study, to obtain a direct association between startle amplitude neural activity startle eyeblink trials from all participants were randomized. Group analyses were conducted on valid trials and invalid trials independently using participants in an emotionally neutral condition. Comparisons between these valid and invalid responses were made regarding underlying neural activity.

## METHODS

### Participants

Seventy-seven white participants (35 males) completed the study for payment. The current analyses an additional 8 participants were excluded due to EEG outliers for startle responses, or if they had fewer than two acceptable startle responses to feedback stimuli.

### Procedure

Participants were seated in front of a computer screen in a soundproofed chamber and were prepared for electroencephalographic and startle recording. Participants were randomly assigned to either a diagnostic math test (DMT) condition, a stressful and emotional context that renders negative goal-relevant stimuli as particularly threatening among women, or a control/problem-solving (PST) condition that is more emotionally neutral and stress free and where goal-relevant stimuli, while still positive or negative valence, is typically perceived as more neutral (i.e., the emotional reaction to positive or negative stimuli is more blunted). In the DMT condition, participants were told that they would be completing tasks that were diagnostic of their math intelligence, while in the PST condition, participants were told that they would be completing tasks that were diagnostic of different types of problem-solving techniques they prefer (Forbes & Leitner, 2014; Forbes et al., 2015). After the instructions were read, participants completed a math feedback task for 34 minutes (described below).

### Pre-Startle Task

Participants completed a 34 minute math task consisting of standard multiplication and division problems (e.g., 10×20=200). This ensured all participants would solve comparable numbers of problems correctly and incorrectly, allowing them to see equal amounts of positive and negative feedback. During each trial, participants were given three answer choices below each problem (A, B or C), with the answer to each problem randomly presented in one of the three answer positions on each trial. Participants were given 16 seconds to solve each problem without scratch paper. Participants made all answer selections via a button box placed in their laps. After each response, participants received feedback for 2 seconds that indicated whether their answer was wrong or correct. If participants were unable to answer a problem within the given 16 seconds they would receive negative (wrong) feedback.

### EEG Recording

Continuous EEG activity was recorded using an ActiveTwo head cap and the ActiveTwo Biosemi system (BioSemi, Amsterdam, Netherlands). Recordings were collected from 128 Ag-AgCl scalp electrodes and from bilateral mastoids. Two electrodes were placed next to each other 1 cm below the right eye to record startle eye-blink responses. A ground electrode was established by BioSemi’s common Mode Sense active electrode and Driven Right Leg passive electrode. EEG activity was digitized with ActiView software (BioSemi) and sampled at 2048 Hz. Data was downsampled post-acquisition and analyzed at 512 Hz.

### Startle Acquisition and extraction

To elicit startle responses, a 40ms burst of white noise (100 db) was presented during the math feedback task through headphones (Bose Quietcomfort 25). During each startle trial, the startle probe elicited one second into the feedback presentation. Startle probes were presented randomly throughout the task. Startle responses were obtained from electromyographic (EMG) recordings of the right orbicularis occuli muscle using two Biosemi FLAT electrodes (BioSemi, Amsterdam, Netherlands). As with the EEG data, startle activity was digitized with ActiView software (Biosemi) and sampled at 2048 Hz. Data was downsampled post-acquisition and analyzed at 512 Hz. Eyeblink EMG data was then filtered by a 28 Hz high-pass FIR filter and then rectified. Peaks appearing between 20ms and 150ms after administration of the acoustic startle probe were operationalized as startle responses (Blumenthal, et al., 2005). Startle trials with eyeblink peak value three standard deviations above the baseline mean (Ornitz, Hanna & de Traversay, 1992) were defined as valid startle response, while trials lower than three SD above the baseline mean were defined as non-response or zero-response trials.

### EEG preprocessing

For startle response analyses EEG signal was epoched and stimulus locked from 500ms pre-feedback presentation to 2000ms post-feedback presentation. EEG artifacts were removed via FASTER (Fully Automated Statistical Thresholding for EEG artifact Rejection; Nolan, et.al., 2010), an automated approach to cleaning EEG data that is based on multiple iterations of independent component and statistical thresholding analyses. In FASTER, EEG trials are down-sampled to 512 Hz, filtered by a 0.3 Hz highpass FIR filter, and baseline corrected using the time series from 100 ms preceding the onset of the tasks. All artifact components are identified and removed as a z-score for that variable, e.g., independent components are identified as eye blink components via their correlation with simultaneous EOG channels and are removed from the EEG signal. The N100 was scored as the average activity at FCz (where it was maximal) between 150-180 ms (when it was peak across all trials). The P200 was scored as the average activity at FCz between 220-260ms, and the P300 was scored as the average activity at CPz between 300-340 ms. EEG trials were recored according its relavant EMG eyeblink peak values, and were grouped as valid startle EEG trials and non-response EEG trials.

### Source reconstruction

All a priori sources used were identified and calculated via forward and inverse models utilized by MNE-python (Gramfort et al., 2013 and Gramfort et al., 2014). The forward model solutions for all source locations located on the cortical sheet were computed using a 3-layers boundary element model (BEM; Hämäläinen and Sarvas, 1989) constrained by the default average template of anatomical MNI MRI. Cortical surfaces extracted with FreeSurfer were sub-sampled to approximately 10,240 equally spaced vertices on each hemisphere. The noise covariance matrix for each individual was estimated from the pre-stimulus EEG recordings after preprocessing. The forward solution matrix and the data were whitened using the noise covariance matrix (Lin et al., 2006). A linear inverse operator was then calculated using the noise covariance matrix and the whitened forward solution, and applied to the EEG data using a loose orientation constraint (loose = 0.2, depth = 0.8; Lin et al., 2006). The current estimates were then normalized to baseline, resulting in a dynamic statistical parametric mapping (dSPM) statistic (Dale et al., 2000), which represents the power of activity within each cortical location (vertex). Then, dSPM values were averaged between certain time period, which is corresponding to specific ERP component, for all vertices for measuring the source-level amplitude. Finally, linear regression was conducted on startle eyeblink amplitude and source activity for every single vertex, an F-value map was generated to represent how closely the brain source activity is related the startle eyeblink amplitude.

## RESULTS

### Distribution of startle elicited eyeblink patterns

The distribution of startle elicited eyeblink patterns is represented in Figure 1. Both the eyeblink peak value distribution (Figure 1a) and distribution of the degree to which eyeblink peak values were larger than baseline (Figure 1b) exhibit an approximate Poisson distribution. Reporting non-response trials varies greatly across studies in terms of how many trials among total trials are labeled as non-response trials, from fewer than 5% (e.g. Meyer et al., 2014) to more than 90% (e.g. Klorman, et al., 2003; Quevedo, et al., 2010). In the present study, 70% of the startle eyeblink trials contained EMG peak values larger than twice the mean of the baseline, and 50% of the trials contained EMG peak values larger than three times the mean of the baseline. Thus, in the current study, half of all startle eyeblink trials did not reach the typical criterion used to label a response trial as above threshold.

**Figure 1.**
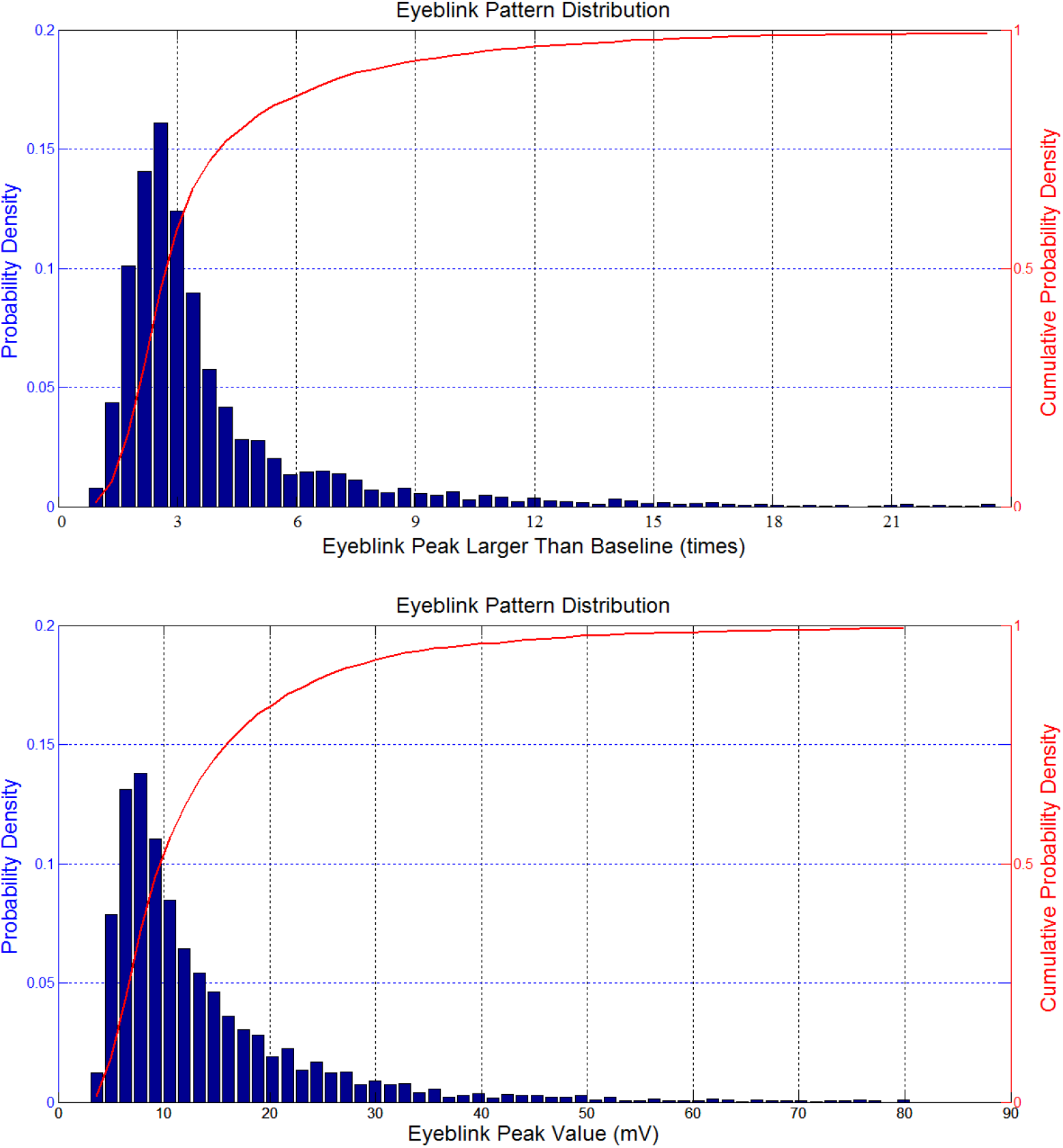
The distribution of startle response eyeblinks, (a) eyeblink peak value distribution, (b) the distribution of the degree to which eyeblink peak value was larger than baseline. Results showed that most of the startle eyeblink trials do not reach the common criterion to be labeled as a response trials.

### ERPs Comparisons for Startle Reflex and Non-Response Startle Reflex ERPs

Based on the sheer number of trials that would normally be excluded it’s imperative to examine whether these exclusions are warranted with respect to the intentions and goals of using the startle probe methodology, which is to measure amygdala activity in response to stimuli of interest. Thus we examined whether there was a difference among ERP waveforms thought to index startle responses, amygdala activity, with respect to trials that would normally be excluded compared to those that would normally be included. To do this we conducted a 2 (feedback type: correct or wrong) × 2 (valid trial: valid or non-response) repeated measures ANOVA on waveform amplitudes for N100, P200 and P300 respectively. Eight participants were removed from this test because they didn’t elicit a startle response belongs to one of the startle type of four. For N100, results revealed a main effect for trial validity, F(1, 67) = 8.625, p = 0.005, where valid startle elicit larger N100 amplitude (M=−2.41, SD=0.45) compared to none-response startle trials (M=−0.91, SD=0.48). No other main effect or interactive effects were found, p’s > 0.234. For P200 and P300, no significant effects were found either in trial validity, feedback type, or interaction between the two (p’s >0.385). Results indicated that participants would release identical startle amplitude regardless of the feedback type they received. Thus, we collapse all correct and wrong feedback startle trials together in following analyses. However, valid and none-response startle trials exhibit significant difference in N100 amplitude, but not in P200 and P300. Figure 2 represents the grand average ERP waveforms of valid (above threshold) and non-response (below threshold) startle reflex. ERPs were generated by averaging startle trials collapsed from all participants.

**Figure 2.**
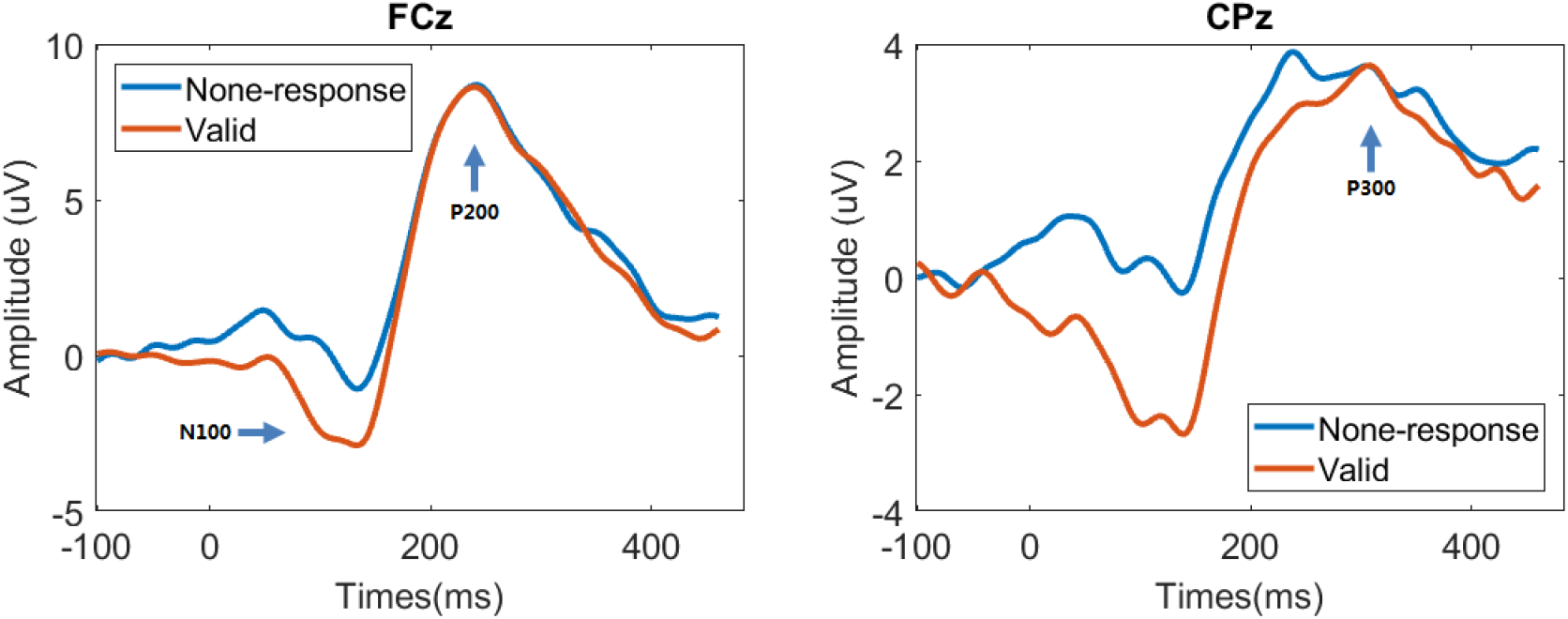
An illustration of ERPs comparison between valid startle trials and non-response startle trials. ERPs were averaged using startle trials collected from all participants. Significant difference in amplitude was found in N100 component between two types of startle trials, but not in P200 and P300 components.

### Trial study on relationship between startle eyeblinks and amplitude of ERPs

We next examined the relationship between startle amplitudes and neural activity in a more sophisticate way, i.e. not only compare below and above threshold startle trials, but separate startle trials in much more levels in terms of their amplitudes. Then we tried to setup relationship between them using linear regression analyses. This also satisfied the increasing demand of analyzing startle fluctuation in dynamic way, in which understanding how startle eyeblink amplitude in relation to neural activity becomes important.

By knowing that valid startle trials have larger N100 amplitude than non-response startle reflex, we next hypothesize that startle eyeblink is associated with N100 amplitude (but not P200 and P300) in general, i.e. below threshold N100 contribute together/equally with above threshold startle N100 generating startle eyeblinks, just in lower amplitude. To test this, we categorized the startle trials EEG into groups in terms of their amplitude, and compare their amplitude with respected to ERP components. However, due to that for each participant, there are too few startle trials to group, we thus gather together all the startle trials regardless of participant, and then make an “*artificial*” ERPs by group the trials depend on their startle eyeblink amplitude *only*. More specifically, 1746 clean startle trials (regardless of their validity and feedback type), were collected across participants. Startle eyeblinks were sorted by amplitude. Every 50 startle trials were grouped together and ERPs were made by averaging the 50 trials of EEG. After sorting and averaging, 35 new trials of startle eyeblinks and ERPs were obtained. All startle trials were separated into a total of 35 groups. Startle eyeblink size was measured by averaging eyeblink peak amplitudes within each group. ERPs were measured by averaging every 50 trials of startle's EEG in each group. Linear regression analyses were conducted on the startle eyeblinks average peak amplitudes and N100, P200 and P300 potentials in each level respectively. Meaningful relationships were found between N100 potentials (p < 0.001) and startle eyeblink sizes. No effects were seen between P200 and P300 potentials (p’s > 0.343) and startle eyeblink size. Complimenting initial ANOVA analyses, these findings suggest that N100 amplitudes exhibited a strong positive relationship with the size of eyeblink peak values (as eyeblink amplitudes increased, N100 amplitudes became more negative), and thus may reflect a linear relationship between amygdala activity and startle responses, regardless of whether the trial would be considered traditionally above or below threshold. These patterns were not reflected in the P200 and P300 components.

### Trial study on relationship between startle eyeblinks and source activity

By knowing that startle eyeblink sizes are strongly correlated to N100 amplitude, regardless of above or below threshold trials. We further hypothesized that brain activity in specific regions, which are supposed to highly interconnected with amygdala, are also strongly associated with startle eyeblink amplitude in N100 time period. This relationship should be also generalized without considering the validity of startle trials. To test this, a data-driven approach was utilized to examine whether there were similar relationships between startle amplitudes and the neural generators of the different components. Finding meaningful linear relationships across trial type would serve as indication that startle amplitudes, regardless of whether they were above threshold, served as a proxy for amygdala activity that varied in intensity. Specifically, linear regression analyses were conducted on brain activity (in dSPM values) elicited in response to startle probes delivered during presentation of feedback. To map a comprehensive representation of the relationship between brain activity and startle eyeblinks, linear regression analyses were conducted on all 20,480 vertices equally distributed on cortical surfaces of each brain. Multiple comparisons/false discovery rates were accounted for by correcting all p-values with Benjamini-Hochberg procedures (Benjamini & Hochberg, 1995). Significant relationship were found using threshold of corrected p-value less than 0.05. Both positive and negative relationships were found. Among the 20,480 vertices, approximately 15% revealed negative relationships with respect to startle eyeblinks. After p-value correction, only small numbers of vertices showed significant negative relationship in all three ERP components. Thus, only positive relationship was represented in our results. Figure 4 displays regions that exhibited significant relationships between brain vertices and startle eyeblinks (corrected p-value<.05).

**Figure 3.**
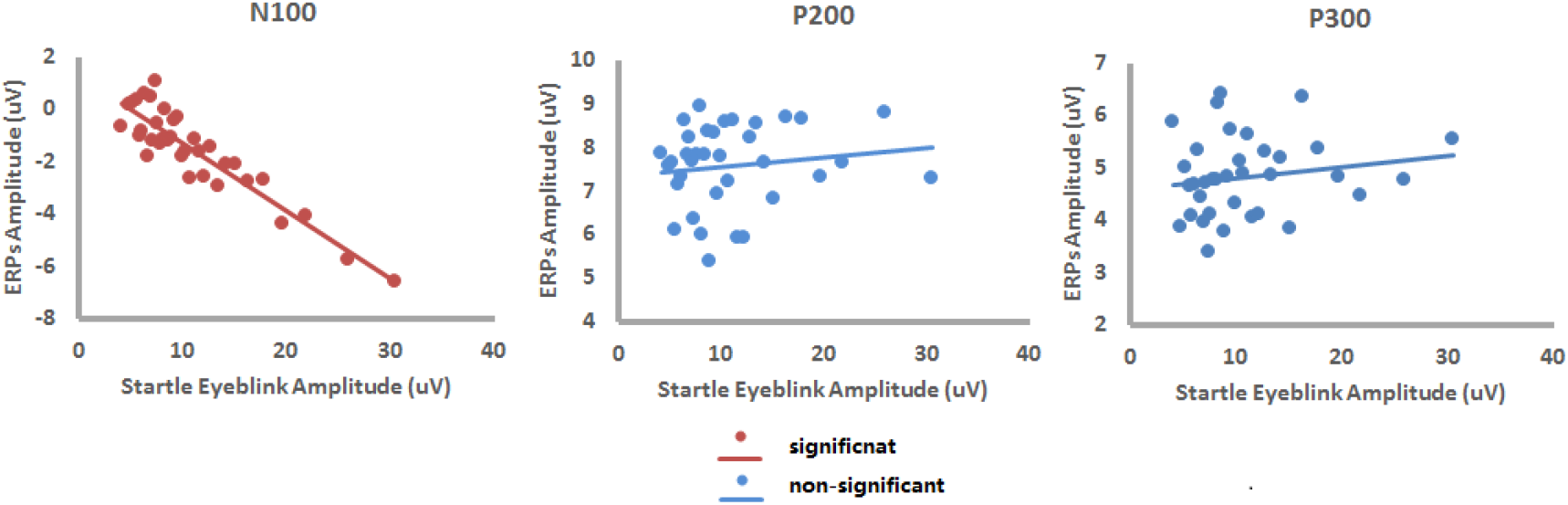
Startle released N100 amplitude exhibits heavily linear relationship with respect to startle eyeblink amplitude (*p<0.001*), however this relationship was not found at all for P200 and P300 (*p’s >0.343*).

**Figure 4.**
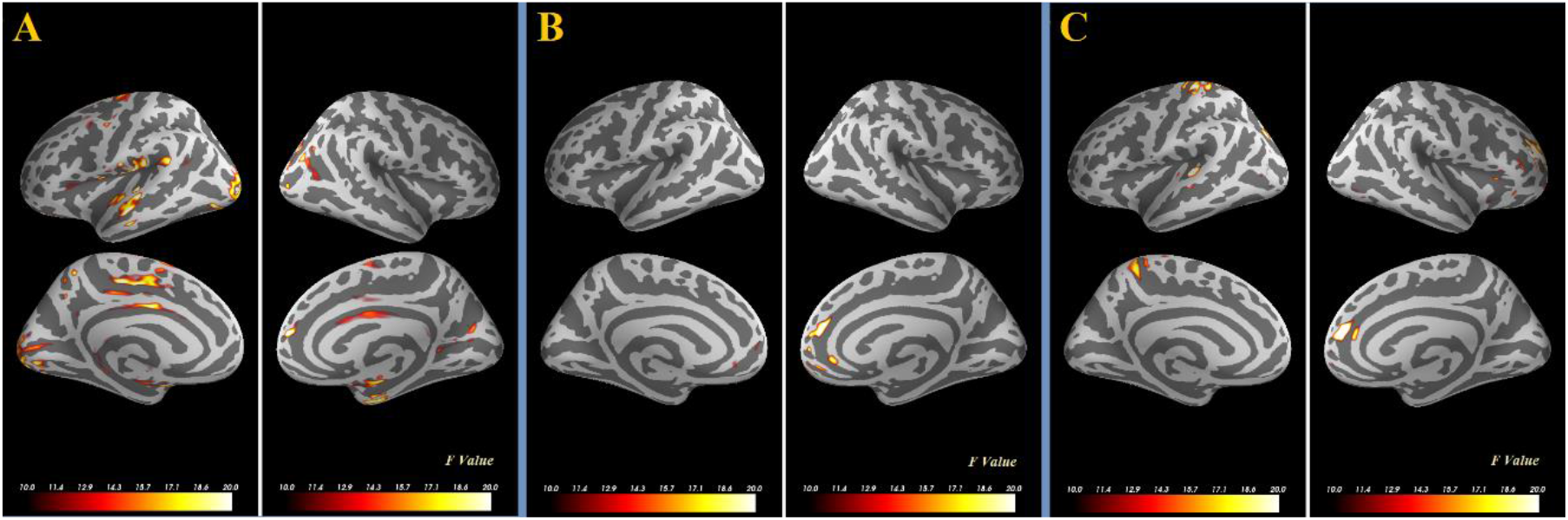
Large amount of brain regions, among which a lot of them are highly interconnected with amygdala, were strongly associated with startle eyeblink amplitude in only N100 time interval, but not in P200 and P300. Results shed light on how brain activity related to startle eyeblink amplitude in twofold: first, even for below threshold startle trials, the brain in large areas may still activated in N100 time interval, but only in a lower manner in amplitude; second, below and above threshold startle trials elicit identical activity in most of the brain regions in P200 and P300 time interval.

**Table 1.** Statistics of the linear regression model on startle eyeblink and the amplitude of ERPs.

|  | R | R<sup>2</sup> | df1 | df2 | F | p |
| --- | --- | --- | --- | --- | --- | --- |
| N100 | 0.913 | 0.835 | 1 | 33 | 166.895 | <0.001 |
| P200 | 0.118 | 0.019 | 1 | 33 | 0.640 | 0.429 |
| P300 | 0.164 | 0.027 | 1 | 33 | 0.927 | 0.343 |

Source analyses demonstrated that within the N100 time interval, brain regions that were strongly associated startle eyeblink size were the primary and secondary auditory cortex, premotor cortex, occipital cortex, dorsal anterior cingulate cortex (dACC), posterior cingulate cortex (PCC), somatosensory cortex, and medial temporal cortex. Activation within the auditory cortex may be related to auditory evoked potentials as the startle response was elicited with a blast of noise (e.g., Papanicolaou et al., 1990). Past research has described activation in the premotor and primary somatosensory cortex have been reported as major contributors in the startle response. Premotor cortex activation has also been elicited as an aversive response to novel and potentially harmful stimuli (Lang et al., 1990) and orientation of attention to the planning of motor execution (Kim et al., 1999; Hari et al., 1982). Relating directly to what the startle response indexes, medial temporal cortex activity has been shown to be directly linked to amygdala activity. This activity has been consistently implicated in perception and response to the danger and threat (LeDoux, 1996; Williams & Gordon, 2007). dACC, PCC and somatosensory cortexes are have also been shown to be highly correlated with amygdala activity. For example, fear potentials reportedly spread to the occipital cortex immediately after the first transient activity elicited in amygdala (Krolak-Salmon et al., 2004). dACC has been shown to be related to detecting impending danger (Eisenberger & Cole, 2012) in addition to modulating danger-related neural networks (Das et al., 2005). PCC may serve an evaluative function for the sensory events (Wyss et al., 2014). Our results indicated that within the time frame of N1, startle reflex strength regulated a large amount of physiological reactions that were widespread in the whole brain. Moreover, large areas of brain regions that are directly correlated to amygdala activity demonstrated the importance of the amygdala in determining the size of the startle eyeblink reflex.

Source analyses demonstrated that within the P200 and P300 time interval, there were also neural regions that exhibited a significant relationship with startle eyeblink amplitude. For example, medial prefrontal cortex (mPFC) and superior frontal cortex (SFC) in P2 and superior frontal extending to ACC, middle frontal gyrus, superior temporal/temporoparietal junction (TPJ), and superior parietal gyrus (SPG) cortices in P3. However, due to the few brain regions found in P2 and P3, this time period may not be as meaningful. In other words, brain activation in P2 and P3 time window does not show meaningful differences with respect to startle eyeblink amplitude.

Overall, results suggest that startle eyeblink amplitude fluctuation only manifests as a fluctuation in brain activity within the N100 time interval, but not P200 and P300. By collapsing all startle trials together, we found that large amount of brain regions, among which a lot of them are highly interconnected with amygdala, were strongly associated with startle eyeblink amplitude. Findings indicate the fluctuation of these large areas of brain regions contributes together to generate the fluctuation of N100. More importantly, results suggest that even for below threshold startle trials, the brain in large areas may still activated, but only in a lower manner in amplitude.

## DISCUSSION

Past research using startle probe methodologies often employ standards that require them to exclude large numbers of trials from their analyses due to factors like habituation and failure to exhibit startle responses among subsets of the population. Yet based on fMRI studies and principles of neural function it stands to reason that amygdala activity, i.e., what startle responses reflect, does not cease to exist on these trials. In this study, neural activity to above and below threshold startle responses were compared with respect to ERPs to obtain a better understanding of how and whether startle responses differ in meaningful ways and/or represent linear aspects of amygdala activation in response to feedback. Results indicated that the N1 component may represent the startle response optimally (i.e., amygdala activity) and to that extent, how above and below threshold startle responses represent linear aspects of the same amygdala response (i.e., less or more amygdala activity). Specifically, when all startle trials from all participants were extracted and grouped according to eyeblink amplitudes, findings indicated that N1 amplitudes predicted startle eyeblink amplitudes, while P2 and P3 amplitudes did not. Source analyses corroborated these findings, demonstrating that within the N1 time interval, source activity in regions typically interconnected with the amygdala and implicated in the startle response exhibited a linear relationship with startle amplitudes. However, during P2 and P3 time intervals, very few brain regions were found to be associated with startle eyeblink amplitude. Given this it can be argued that brain activity during the N1 time interval most likely reflects amygdala activity and that both above and below threshold startle responses reflect amygdala activity as well.

Importantly, our findings suggest that excluding below threshold startle trials could ultimately mask meaningful patterns and fluctuations in amygdala activity otherwise present in a given experiment. While past research utilizing startle probe methodologies found marked linear decreases in amygdala responses to stimuli over time, i.e., a habituation effect, past research in fMRI has revealed meaningful non-linear activations in the amygdala that fluctuates over time in response to positive and negatively valanced stimuli (Phillips et al., 2001). We have uncovered similar patterns as well by including below threshold trials in some of our past work.

For instance, Forbes et al. (2018) exposed men and women to a manipulation that made the situation particularly stressful for women and rendered negative (compared to positive) feedback received in these contexts as threatening. When below threshold startle trials were excluded, and several subjects were excluded for not having sufficient numbers of above threshold startle trials, analyses revealed a linear relationship indicating that all subjects elicited larger startle responses to negative feedback, and women elicited larger startle responses in general compared to men. When including below threshold trials, however, quadratic over time models indicated that startle amplitudes significantly fluctuated only for women in the stressful condition and in response to negative feedback specifically, which was hypothesized to be more threatening in these contexts (and thus more likely to evoke amygdala activity). Inclusion of the below threshold startle trials thus yielded a much more nuanced understanding of the complex relationship between the amygdala and stressful experiences in specific contexts that otherwise are not evident when analyses excluded below threshold trials.

Source and linear regression analyses conducted on the N1 time interval not only revealed linear relationships between activity in a given region and above and below threshold startle trials, but they yielded activity in regions heavily interconnected with the amygdala, and regions thought to be a part of a broader, amygdala centric network involved in attentional reorientation and allocation and arousal in general. This can be further verified by connectivity-based analysis (Liu et al., 2011; Liu et al., 2012; Liu et al., 2013). Specifically, past literature have shown that amygdala activity modulates in the large amount of brain regions, e.g. auditory cortex, occipital cortex, premotor cortex hippocampus, anterior cingulate cortex (ACC), and posterior cingulate cortex (PCC), modulating the startle reflex accordingly (Medford & Nick, 2010; Lee, Younglim, 1997, Cristina, et al., 2009). Bolstering this rich literature, in this study regression analyses yielded significant activity within primary and secondary auditory cortex, premotor cortex, occipital cortex, dorsal anterior cingulate cortex (dACC), posterior cingulate cortex (PCC), somatosensory cortex, and medial temporal cortex.

Amygdala activity has been shown to orchestrate whole brain dynamics in the presence of a threat, such as a startle probe, as it is often described as a regulator of the stress response and a hub for emotional processing (Liu et al., 2017). For example, medial temporal cortex activity has been shown to be directly linked to amygdala activity (Liu et al., 2015; Amey et al., 2018). This activity has been consistently implicated in perception and response to the danger and threat (LeDoux, 1996; Williams & Gordon, 2007; Eldawud et al., 2016). dACC, PCC and somatosensory cortexes have also been shown to be highly correlated with amygdala activity. For example, fear potentials reportedly spread to the occipital cortex immediately after the first transient activity elicited in amygdala (Krolak-Salmon et al., 2004). dACC has been shown to be related to detecting impending danger (Eisenberger & Cole, 2012) in addition to modulating danger-related neural networks (Das et al., 2005). In sum, the downstream activity of amygdala activation orchestrates activity of the hippocampus, and areas in the cortex, including dorsal and ventral aspects of anterior cingulate and prefrontal cortex, bilateral insular cortex and bilateral middle frontal gyri (LaBar & Cabeza, 2006; Bush, Luu, & Posner, 2000; Kim, 2011; Murty, Ritchey, Adcock, & LaBar, 2010; Zald et al., 1998; He et al., 2019). When the present analyses focused on the N100 time period, results supported these past conjectures, in addition to expanding on them as we were able to index this deep brain activity by using regions closer to the scalp as a proxy. Results indicated that within the time frame of N1, startle reflex strength regulated a large amount of physiological reactions that were widespread in the whole brain. However, considering the significant involvement of the amygdala in startle and emotional responses in very early stage of startle reflex (LeDoux, 1996; Alessandro, et cl. 1996), suggesting that in N1 time interval, startle eyeblink size may represent amygdala activity rather than any other cognitive processes.

In the current study, for P200 and P300, there were no meaningful differences between amplitudes or neural generator activation between these two waveforms with respect to startle eyeblink amplitude. In past literatures, P300 was thought to index attentional and initial memory storage events (Polich, 2007), and P2 has been previously associated with later emotional cognitive processes (Somme, Molen & Pascalis, 2016) and sensory motor gating (Boutros, et.cl., 1999; Pascalis, Cozzuto & Russo, 2013; Zhu et al., 2019). Our results might suggest that regardless of the initial response to the startle stimulus, individuals may still elicit enough attention to the stimulus presented. Habituation is usually found in startle eyeblink tracking; thus, these habituated trials are often excluded. The finding in present study also illustrates that below threshold startle reflexes may only represent devaluation in terms of emotional or sensory reaction - these trials, on the contrary, result in almost identical attentional arousal level in comparison to above threshold startle trials.

In this study, regression analysis was conducted on startle trials collected from multiple participants, i.e. ERPs were created by averaging trials with similar startle eyeblink sizes, but not by averaging trials extracted from same individual. In this case, the amplitude of ERPs indexes only direct physiological association with startle eyeblink amplitude. Individual differences, context, and environment noise were not considered. Future studies will work on the role individual differences play in tracking startle eyeblink amplitudes and their relationships with cognitive functions. This study contained many non-response startle trials (about 50%), more than usually reported in past literature. One possible reason may be that solving complex math problems before receiving startle stimulus may occupied participants working memory, decreasing attentional resources. This decrease in attentional resources may have reduced the startle reflex eyeblink sizes.

Although thousands of trials were used in our analyses, providing a means to isolate deeper signals in the brain, it is also always important with respect to EEG to stress caution in interpretation of results related to specific brain regions given the spatial limitations inherent in the methodology. Nevertheless, by using a high-density electrode array, employing an advanced source bayesian analytic approach including dSPM inverse operators, and confining analyses to regions closer to the cortical surface, it is possible to make accurate assumptions about contributions from specific brain regions (Cohen, 2014).

Using a temporal-spatial data-driven approach (Mantena et al., 2009), results indicate fluctuating startle eyeblink amplitude could approximate fluctuating amygdala activity. Previously, startle eyeblink amplitude has had a limited role in measuring fluctuating activity in the amygdala. Findings suggest that non-response startle eyeblink amplitudes can provide meaningful data, as non-response trials were shown to exhibit meaningful patterns within the N100 time period, equivalent to that of valid response trials. Although detailed mechanisms of startle eyeblink fluctuation are still unclear, these attention and memory process within self-threatening contexts, e.g. stressful situation in this study, may cause later increases in amygdala activity, and consequently startle responses. Future studies could be placed on how these ERP components may contribute to this effect. Results shed light on why it is still important to include these non-response startle trials in analyses providing evidence from both neural signatures and over time analyses. Valuable contributions to this question can be answered by future work describing how low amplitude non-startle response trials can be affected by cognitive processing and provide insight into underlying amygdala activity.

## ACKNOWLEDGEMENTS

All aspects of this study and article were supported by National Science Foundation grants #1329281 and #1535414 awarded to Chad Forbes.

## REFERENCE

Amey, R., Leitner, J., Liu, M., & Forbes, C. (2018). Neural Mechanisms Associated with Semantic and Basic Self-Oriented Memory Processes Interact to Modulate Self-Esteem. bioRxiv, 350926. DOI: https://doi.org/10.1101/350926

Antonova, E., Chadwick, P., & Kumari, V. (2015). More meditation, less habituation? The effect of mindfulness practice on the acoustic startle reflex.PloS one, 10(5), e0123512.

Armstrong, B. C., Ruiz-Blondet, M. V., Khalifian, N., Kurtz, K. J., Jin, Z., & Laszlo, S. (2015). Brainprint: Assessing the uniqueness, collectability, and permanence of a novel method for ERP biometrics. Neurocomputing, 166, 59–67.

Bradley, M. M., Codispoti, M., & Lang, P. J. (2006). A multi-process account of startle modulation during affective perception. Psychophysiology,43(5), 486–497.

Bradley, M. M., Lang, P. J., & Cuthbert, B. N. (1993). Emotion, novelty, and the startle reflex: habituation in humans. Behavioral neuroscience, 107(6), 970.

Braff, D. L., Geyer, M. A., & Swerdlow, N. R. (2001). Human studies of prepulse inhibition of startle: normal subjects, patient groups, and pharmacological studies. Psychopharmacology, 156(2-3), 234–258.

Blumenthal, T. D., Cuthbert, B. N., Filion, D. L., Hackley, S., Lipp, O. V., & Van Boxtel, A. (2005). Committee report: Guidelines for human startle eyeblink electromyographic studies. Psychophysiology, 42(1), 1–15.

Brinkworth, R.S.A. &Turker, K. S. (2003). A method for quantifying reflex responses from intra-muscular and surface electromyogram. Journal of Neuroscience Methods, 122, 179–193

Cuthbert, B. N., Schupp, H. T., Bradley, M., McManis, M., & Lang, P. J. (1998). Probing affective pictures: Attended startle and tone probes.Psychophysiology, 35(03), 344–347.

Das, P., Kemp, A. H., Liddell, B. J., Brown, K. J., Olivieri, G., Peduto, A., … & Williams, L. M. (2005). Pathways for fear perception: modulation of amygdala activity by thalamo-cortical systems. Neuroimage, 26(1), 141–148.

De Pascalis, V., Cozzuto, G., & Russo, E. (2013). Effects of personality trait emotionality on acoustic startle response and prepulse inhibition including N100 and P200 event-related potential. Clinical Neurophysiology, 124(2), 292–305.

De Pascalis, V., & Russo, E. (2013). Hypnotizability, hypnosis and prepulse inhibition of the startle reflex in healthy women: An ERP analysis. PloS one,8(11), e79605.

Delplanque, S., Lavoie, M. E., Hot, P., Silvert, L., & Sequeira, H. (2004). Modulation of cognitive processing by emotional valence studied through event-related potentials in humans. Neuroscience letters, 356(1), 1–4.

Delplanque, S., Silvert, L., Hot, P., & Sequeira, H. (2005). Event-related P3a and P3b in response to unpredictable emotional stimuli. Biological psychology, 68(2), 107–120.

Dien, J., Spencer, K. M., & Donchin, E. (2003). Localization of the event-related potential novelty response as defined by principal components analysis. Cognitive Brain Research, 17(3), 637–650.

Eisenberger, N. I., & Cole, S. W. (2012). Social neuroscience and health: neurophysiological mechanisms linking social ties with physical health.Nature neuroscience, 15(5), 669–674.

Eldawud, R., Reitzig, M., Opitz, J., Rojansakul, Y., Jiang, W., Nangia, S., & Dinu, C. Z. (2016). Combinatorial approaches to evaluate nanodiamond uptake and induced cellular fate. Nanotechnology, 27(8), 085107.

Engelmann, J. M., Gewirtz, J. C., & Cuthbert, B. N. (2011). Emotional reactivity to emotional and smoking cues during smoking abstinence: Potentiated startle and P300 suppression. Psychophysiology, 48(12), 1656–1668.

Falls, W. A., Miserendino, M. J., & Davis, M. (1992). Extinction of fear-potentiated startle: blockade by infusion of an NMDA antagonist into the amygdala. Journal of Neuroscience, 12(3), 854–863.

Filion, D. L., Dawson, M. E., & Schell, A. M. (1998). The psychological significance of human startle eyeblink modification: a review. Biological psychology, 47(1), 1–43.

Forbes, C. E. (2015). On social neuroscience methodologies and their applicability to group processes and intergroup relations. Group Processes & Intergroup Relations, 18(3), 348–365.

Forbes, C. E., Amey, R., Magerman, A. B., Duran, K., & Liu, M. (2018). Stereotype-based stressors facilitate emotional memory neural network connectivity and encoding of negative information to degrade math self-perceptions among women. Social cognitive and affective neuroscience, 13(7), 719–740.

Forbes, E. E., Miller, A., Cohn, J. F., Fox, N. A., & Kovacs, M. (2005). Affect-modulated startle in adults with childhood-onset depression: Relations to bipolar course and number of lifetime depressive episodes. Psychiatry Research, 134(1), 11–25.

Forbes, C. E., & Leitner, J. B. (2014). Stereotype threat engenders neural attentional bias toward negative feedback to undermine performance. Biological psychology, 102, 98–107.

Gard, M. G., & Kring, A. M. (2007). Sex differences in the time course of emotion. Emotion, 7(2), 429.

Grillon, C., Baas, J. P., Lissek, S., Smith, K., & Milstein, J. (2004). Anxious responses to predictable and unpredictable aversive events. Behavioral neuroscience, 118(5), 916.

Gold, A. L., Morey, R. A., & McCarthy, G. (2015). Amygdala–prefrontal cortex functional connectivity during threat-induced anxiety and goal distraction. Biological psychiatry, 77(4), 394–403.

Hajcak, G., & Foti, D. (2008). Errors are aversive defensive motivation and the error-related negativity. Psychological Science, 19(2), 103–108.

Hari, R., Kaila, K., Katila, T., Tuomisto, T., & Varpula, T. (1982). Interstimulus interval dependence of the auditory vertex response and its magnetic counterpart: implications for their neural generation. Electroencephalography and clinical neurophysiology, 54(5), 561–569.

He, X., Zhu, Y., Lin, Y. C., Li, M., Du, J., Dong, H., … & Zhang, L. (2019). PRMT1-mediated FLT3 arginine methylation promotes maintenance of FLT3-ITD+ Acute Myeloid Leukemia. Blood, blood–2019001282.

Herbert, C., Kissler, J., Junghöfer, M., Peyk, P., & Rockstroh, B. (2006). Processing of emotional adjectives: Evidence from startle EMG and ERPs.Psychophysiology, 43(2), 197–206.

Herten N, Pomrehn D, Wolf OT. Memory for objects and startle responsivity in the immediate aftermath of exposure to the Trier Social Stress Test. Behavioural brain research. 2017 May 30;326:272–80.

Hillman, C. H., Hsiao-Wecksler, E. T., & Rosengren, K. S. (2005). Postural and eye-blink indices of the defensive startle reflex. International Journal of Psychophysiology, 55(1), 45–49.

Jiang, W., Luo, J., & Nangia, S. (2015). Multiscale approach to investigate self-assembly of telodendrimer based nanocarriers for anticancer drug delivery. Langmuir, 31(14), 4270–4280.

Jiang, W., Wang, X., Guo, D., Luo, J., & Nangia, S. (2016). Drug-specific design of telodendrimer architecture for effective doxorubicin encapsulation. The Journal of Physical Chemistry B, 120(36), 9766–9777.

Karla, S., Ruusuvirta, T., & Wikgren, J. (2014). Effect of Emotional Picture Viewing on Voluntary Eyeblinks. PloS one, 9(3), e89536.

Keil, A., Bradley, M. M., Junghöfer, M., Russmann, T., Lowenthal, W., & Lang, P. J. (2007). Cross-modal attention capture by affective stimuli: evidence from event-related potentials. Cognitive, Affective, & Behavioral Neuroscience, 7(1), 18–24.

Key, A. P. F., Dove, G. O., & Maguire, M. J. (2005). Linking brainwaves to the brain: an ERP primer. Developmental neuropsychology, 27(2), 183–215.

Klorman, R., Cicchetti, D., Thatcher, J. E., & Ison, J. R. (2003). Acoustic startle in maltreated children. Journal of Abnormal Child Psychology, 31(4), 359–370.

Krolak-Salmon, P., Hénaff, M. A., Vighetto, A., Bertrand, O., & Mauguière, F. (2004). Early amygdala reaction to fear spreading in occipital, temporal, and frontal cortex: a depth electrode ERP study in human. Neuron, 42(4), 665–676.

Lang, P. J. (1995). The emotion probe: Studies of motivation and attention. American psychologist, 50(5), 372.

Lang, P. J., Bradley, M. M., & Cuthbert, B. N. (1990). Emotion, attention, and the startle reflex. Psychological review, 97(3), 377.

Lang A, Newhagen J, Reeves B. Negative video as structure: Emotion, attention, capacity, and memory. Journal of Broadcasting & Electronic Media. 1996 Sep 1;40(4):460–77.

Leite, J., Carvalho, S., Galdo-Alvarez, S., Alves, J., Sampaio, A., & Gonçalves, Ó. F. (2012). Affective picture modulation: Valence, arousal, attention allocation and motivational significance. International Journal of Psychophysiology, 83(3), 375–381.

Lissek, S., Baas, J. M., Pine, D. S., Orme, K., Dvir, S., Nugent, M., … & Grillon, C. (2005). Airpuff startle probes: an efficacious and less aversive alternative to white-noise. Biological psychology, 68(3), 283–297.

Liu, M., Amey, R. C., & Forbes, C. E. (2017). On the role of situational stressors in the disruption of global neural network stability during problem solving. Journal of cognitive neuroscience, 29(12), 2037–2053.

Liu, M., Kuo, C. C., & Chiu, A. W. (2011). Statistical threshold for nonlinear granger causality in motor intention analysis. In 2011 Annual International Conference of the IEEE Engineering in Medicine and Biology Society (pp. 5036–5039). IEEE.

Liu, M. T., Kuo, C. C., & Chiu, A. W. (2013). Non-linear Granger causality and its frequency decomposition in decoding human upper limb movement intentions. International Journal of Biomedical Engineering and Technology 34, 12(1), 1–25.

Liu, M., Forbes, C., Jordan, K., & Magerman, A. (2015, September). UNIQUE SPATIAL AND SPECTRAL EEG BIOMARKERS CHARACTERIZE THE EMG STARTLE RESPONSE. In PSYCHOPHYSIOLOGY (Vol. 52, pp. S25–S25). 111 RIVER ST, HOBOKEN 07030-5774, NJ USA: WILEY-BLACKWELL.

Liu, M., & Wang, X. (2017). Beyond the ERPs—Startle Response is Better Outlined by Whole Brain and Spectral EEG Features. Journal of Psychiatry and Brain Science, 2(3).

Mangun, G. R. (1995). Neural mechanisms of visual selective attention. Psychophysiology, 32(1), 4–18.

Mantena, V., Jiang, W., Li, J., & McKenzie, R. (2009, April). Prostate cancer biomarker identification using MALDI-MS data: initial results. In 2009 IEEE/NIH Life Science Systems and Applications Workshop (pp. 116–119). IEEE.

Marinkovic, K., Halgren, E., & Maltzman, I. (2001). Arousal-related P3a to novel auditory stimuli is abolished by a moderately low alcohol dose. Alcohol and Alcoholism, 36(6), 529–539.

Meyer, T., Quaedflieg, C. W., Giesbrecht, T., Meijer, E. H., Abiad, S., & Smeets, T. (2014). Frontal EEG asymmetry as predictor of physiological responses to aversive memories. Psychophysiology, 51(9), 853–865.

Miller, M. W., Patrick, C. J., & Levenston, G. K. (2002). Affective imagery and the startle response: Probing mechanisms of modulation during pleasant scenes, personal experiences, and discrete negative emotions.Psychophysiology, 39(4), 519–529.

Nelson B.D., Hajcak G. Defensive motivation and attention in anticipation of different types of predictable and unpredictable threat: A startle and event-related potential investigation. Psychophysiology. 2017 Aug 1;54(8):1180–94.

Nelson, B. D., Hajcak, G., & Shankman, S. A. (2015). Event-related potentials to acoustic startle probes during the anticipation of predictable and unpredictable threat. Psychophysiology, 52(7), 887–894.

Nolan, H., Whelan, R., & Reilly, R. B. (2010). FASTER: fully automated statistical thresholding for EEG artifact rejection. Journal of neuroscience methods, 192(1), 152–162.

Olofsson, J. K., Nordin, S., Sequeira, H., & Polich, J. (2008). Affective picture processing: an integrative review of ERP findings. Biological psychology, 77(3), 247–265.

Panayiotou, G., van Oyen Witvliet, C., Robinson, J. D., & Vrana, S. R. (2011). A startling absence of emotion effects: Active attention to the startle probe as a motor task cue appears to eliminate modulation of the startle reflex by valence and arousal. Biological psychology, 87(2), 226–233.

Papanicolaou, A. C., Baumann, S., Rogers, R. L., Saydjari, C., Amparo, E. G., & Eisenberg, H. M. (1990). Localization of auditory response sources using magnetoencephalography and magnetic resonance imaging. Archives of Neurology, 47(1), 33–37.

Picton, T. W., Goodman, W. S., & Bryce, D. P. (1970). Amplitude of evoked responses to tones of high intensity. Acta oto-laryngologica, 70(2), 77–82.

Phillips, M. L., Medford, N., Young, A. W., Williams, L., Williams, S. C., Bullmore, E. T., … & Brammer, M. J. (2001). Time courses of left and right amygdala responses to fearful facial expressions. Human brain mapping, 12(4), 193–202.

Polich, J. (2003). Theoretical overview of P3a and P3b. In Detection of Change (pp. 83–98). Springer US.

Polich, J. (2007). Updating P300: an integrative theory of P3a and P3b.Clinical neurophysiology, 118(10), 2128–2148.

Poli, E., & Angrilli, A. (2015). Greater general startle reflex is associated with greater anxiety levels: a correlational study on 111 young women. Frontiers in behavioral neuroscience, 9, 10.

Putnam, L. E., & Roth, W. T. (1990). Effects of stimulus repetition, duration, and rise time on startle blink and automatically elicited P300.Psychophysiology, 27(3), 275–297

Quevedo, K., Smith, T., Donzella, B., Schunk, E., & Gunnar, M. (2010). The startle response: Developmental effects and a paradigm for children and adults. Developmental psychobiology, 52(1), 78–89.

Ramirez-Moreno, D. F., & Sejnowski, T. J. (2012). A computational model for the modulation of the prepulse inhibition of the acoustic startle reflex.Biological cybernetics, 106(3), 169–176.

Rhudy, J. L., Williams, A. E., McCabe, K. M., Russell, J. L., & Maynard, L. J. (2008). Emotional control of nociceptive reactions (ECON): do affective valence and arousal play a role?. Pain, 136(3), 250–261.

Robinson, J. D., & Vrana, S. R. (2000). The time course of emotional and attentional modulation of the startle eyeblink reflex during imagery.International Journal of Psychophysiology, 37(3), 275–289.

Roth, W. T., Dorato, K. H., & Kopell, B. S. (1984). Intensity and task effects on evoked physiological responses to noise bursts. Psychophysiology,21(4), 466–481.

Sabatinelli, D., Bradley, M. M., & Lang, P. J. (2001). Affective startle modulation in anticipation and perception. Psychophysiology, 38(04), 719–722.

Schupp, H. T., Cuthbert, B. N., Bradley, M. M., Birbaumer, N., & Lang, P. J. (1997). Probe P3 and blinks: Two measures of affective startle modulation.Psychophysiology, 34(1), 1–6.

Sommer, K., van der Molen, M. W., & De Pascalis, V. (2016). BIS/BAS sensitivity and emotional modulation in a prepulse-inhibition paradigm: A brain potential study. Physiology & behavior, 154, 100–113.

Sugawara, M., Sadeghpour, M., De Traversay, J., & Ornitz, E. M. (1994). Prestimulation-induced modulation of the P300 component of event related potentials accompanying startle in children. Electroencephalography and clinical neurophysiology, 90(3), 201–213.

Townsend, J. D., Torrisi, S. J., Lieberman, M. D., Sugar, C. A., Bookheimer, S. Y., & Altshuler, L. L. (2013). Frontal-amygdala connectivity alterations during emotion downregulation in bipolar I disorder. Biological psychiatry,73(2), 127–135.

Tremblay, K., Kraus, N., McGee, T., Ponton, C., & Otis, B. (2001). Central auditory plasticity: changes in the N1-P2 complex after speech-sound training. Ear and hearing, 22(2), 79–90.

Walker, D. L., & Davis, M. (2002). The role of amygdala glutamate receptors in fear learning, fear-potentiated startle, and extinction. Pharmacology Biochemistry and Behavior, 71(3), 379–392.

Wunderlich, J. L., & Cone-Wesson, B. K. (2001). Effects of stimulus frequency and complexity on the mismatch negativity and other components of the cortical auditory-evoked potential. The Journal of the Acoustical Society of America, 109(4), 1526–1537.

Yeomans, J. S., & Frankland, P. W. (1995). The acoustic startle reflex: neurons and connections. Brain research reviews, 21(3), 301–314.

Zhu, Y., He, X., Lin, Y. C., Dong, H., Zhang, L., Chen, X., … & Sun, J. (2019). Targeting PRMT1-mediated FLT3 methylation disrupts maintenance of MLL-rearranged acute lymphoblastic leukemia. Blood, 134(15), 1257–1268.

